# A Clinically Aligned Murine Model of Electroconvulsive Stimulation Reverses Social Aversion and Displays Fear Memory Impairment After Chronic Social Defeat Stress

**DOI:** 10.1101/2025.02.12.637719

**Authors:** Michael D. Kritzer, Stephanie A. Maddox, Michelle X. Chen, Shu Dan, Herbert E. Covington, Klaus Miczek, Joan A. Camprodon, Torsten Klengel, Kerry J. Ressler

## Abstract

Electroconvulsive therapy (ECT) is the most effective treatment for patients with major depression, bipolar depression, mania, catatonia, and schizophrenia. Nonetheless, the mechanisms underlying its therapeutic effects largely remain unknown. While previous preclinical studies have noted a role for neurotropic signaling, neurogenesis, and alterations in monoamine neurotransmitter systems, these models were largely conducted using procedures that deviate from clinical practice. Therefore, we sought to develop a clinically relevant murine model of ECT, referred to as electroconvulsive stimulation (ECS) in animals, which more closely aligns with current clinical approaches to better explore its mechanisms. Using the well-established chronic social defeat stress (CSDS) paradigm, known to negatively impact reward processes, we investigated whether the behavioral changes after CSDS could be reversed following a clinically related course of ECS. Additionally, we observed induction of plasticity-related genes in the nucleus accumbens (NAcc) and amygdala, regions responsible for reward and fear-related memory, respectively. Lastly, we investigated ECS-related changes in the NAcc with bulk RNA-sequencing. Pathway analysis demonstrated cellular changes primarily involved in neuroplasticity and regulating cell migration and differentiation. Therefore, utilizing our novel and clinically relevant model of ECS, we have begun to elucidate mechanisms that contribute to ECT’s therapeutic outcomes by examining murine behavior and RNA from brain regions associated with stress-induced states that model anxiety and depression and the effects of ECS.

## Introduction

Major depressive disorder (MDD) is the most common mental illness and one of the most disabling diseases, affecting more than 300 million people worldwide (World Health Organization, 2023). It is a brain-wide syndrome that often manifests as symptoms of low mood, anhedonia, amotivation, rumination, dysregulated sleep, appetite, cognition, and suicidal thoughts. While there are effective treatments, if first-line approaches fail, their efficacy diminishes with each additional attempt (Rush AJ, 2006). Treatment-resistant depression (TRD), defined as failing two or more antidepressants within a given depressive episode, comes with a high burden of personal and societal cost. Used for over 80 years, a series of clinically induced seizures called electroconvulsive therapy (ECT), remains most effective treatment for TRD; however it is accompanied by general anesthesia, stigma, and cognitive side effects that limit its use (Nuninga JO, 2018). Though there have been advances in somatic therapies, there remains a need to develop novel treatments that work when others fail.

ECT has evolved since the 1940’s and is now routinely used to treat depression, mania, catatonia, and schizophrenia. Little is known about ECTs mechanisms of efficacy and cognitive impairment. ECT induces many seemingly nonspecific neuroplastic changes, making it challenging to interpret. Clinical studies have found whole brain increases in gray matter (Sartorius A D. T.-J., 2018), (Ota M, 2015). One common observation, as seen with other antidepressant treatments, is an increase in hippocampal gray matter following ECT (Sartorius A D. T., 2015), though it is still not clear whether the observed changes in human neuroimaging mediate its therapeutic effects. While ECT is non-focal, increasing evidence points towards a fundamental role in modulating the mesocorticolimbic reward circuitry. The mesocorticolimbic system is a complex multifunctional network responsive to positive and negative reinforcement essential for emotional, cognitive, and motivational abnormalities in depression (Nestler & Carlezon, 2006). The core regions of this reward circuitry are a complex network of richly interconnected axonal pathways that include the hippocampus, nucleus accumbens (NAcc), prefrontal cortex (PFC), cingulate cortex, amygdala, and frontostriatal connections (Russo SJ, 2013).

Human neuroimaging has provided evidence for ECT’s circuit-specific effects that mediate reward via its clinical dimensions: anhedonia and amotivation. Dysregulation of the NAcc and its connections have been associated with increased anhedonia observed in depression (Naranjo CA, 2001). Using multimodal MRI to study patients with depression undergoing ECT, we have observed increased volume in the VTA and NAcc that correlate with clinical improvement (unpublished findings). In addition, diffusion tensor imaging (DTI) identified the mesocorticolimbic fibers connecting the VTA to the NAcc and the cortex and identified tract-specific changes in functional anisotropy that also correlated with improvement in human anhedonia. A similar mesolimbic pathway consisting of VTA dopaminergic projections to the NAcc contribute to the rodent reward system and have been shown to mediate motivation for reward. Not being able to determine causal and molecular mechanisms of psychiatric disorders and their treatments is a major drawback of human imaging studies that has led to preclinical work with rodent electroconvulsive stimulation (ECS), the animal model of ECT, for investigating fundamental mechanisms.

The amygdala is crucial to emotional learning and results in rapid responses to similar threats in the future (Drevets WC, 1992). Fear learning and extinction are processes that involve the amygdala (JE, 1993) and are dysregulated in neuropsychiatric disorders. In humans, fMRI studies have shown amygdala activation in response to fearful visual stimuli (Phelps EA, 2005), consistent with findings across species of amygdala hyperactivation with exposure fearful situations. Stressors like bullying, abuse, and trauma, activate this circuit and can have maladaptive consequences resulting in trauma-spectrum disorders including post-traumatic stress disorder (PTSD), anxiety, and mood disorders. Although hippocampal fMRI studies have been generally consistent in showing volumetric decreases in patients with MDD, similar fMRI studies in the amygdala have been inconsistent (Frodl T, 2006), (Raffaele Cacciaglia, 2018). These inconsistencies could be because the amygdala can be subdivided and different neural circuits extending from its different regions (Maddox SA, 2019). Despite these inconsistencies, the hyperactivity of the amygdala in mood and anxiety disorders has been well established.

Exposure to stress promotes susceptibility to depression (Duman, 2016). The chronic social defeat stress (CSDS) paradigm is an established model for understanding depression mechanisms (Kudriavtseva NN, 1989). The model utilizes social conflict and involves a larger, resident mouse attacking (Sugama S, 2016) a smaller, mouse after its placed within the larger mouse’s territory. Physiological changes consistent with human depression occur with CSDS, such as altered circadian rhythms, increased heart rate, and changes in core temperature (Tornatzky W, 1993). CSDS also has a wide range of behavioral effects, which have relevance to depression mechanisms. It has been shown to increase immobility in the forced swim test (Bondar N, 2018), induce social avoidance (Krishnan V, 2007), (Berton O, 2006), decrease sucrose preference, and increase time spent in the closed arms of the elevated plus maze (Venzala E, 2012). Social contact is rewarding in mice (Panksepp JB, 2007) and social isolation is also common individuals with depression (CB, 1973). CSDS has a range of neurobiological effects in conjunction with the behavioral alterations. It has been shown to decrease the size of the cingulate cortex, NAcc, thalamus, and bed nucleus of the stria terminalis (BNST) and increase the volume of the VTA, hypothalamus, and amygdala by altering the spine density in mice (Anacker C, 2016). Volumetric changes of the medial PFC also appear to be reversible with antidepressant treatment (Czéh B M. T., 2001), (Czéh B L. P., 2007). Specifically, the VTA has been shown to be involved in social avoidance after CSDS (Krishnan V, 2007), yet it is unknown whether the NAcc is directly involved in VTA social processing. The NAcc has been shown to be involved in social reward learning in mice as well as anhedonia in humans (Liu R, 2021), (He ZX, 2020). Clinical approaches that encourage social interaction improves depression and decreases the risk of suicide (Szanto K, 2021), (Kleiman EM, 2013), (Elmer, 2020). Social reward and aversion are driven primarily by the mesolimbic system (Alcaro A, 2007), (Yuan L, 2019).

Behaviorally, chronic ECS has been shown to decrease immobility during the forced swim test (Bingjin Li, 2007), but impairs memory in spatial mazes and in conditioned emotional response, a test of emotional learning (C. Kyeremanteng, 2014). This impaired aversion memory may help model the negative cognitive effects on episodic memory produced by ECT in humans (Fraser LM, 2008). This suggests that ECS is a relevant murine model of ECT in reversing behavioral effects after chronic stress. Additional work is needed to determine whether ECS can model molecular and circuit effects. Appropriate clinical translation of ECT will provide more accurate insights when studying its mechanisms. Furthermore, previous models of ECS occurred under the absence of clinically relevant anesthesia, which is known to alter neuronal function, i.e., seizure threshold (Bundy, 2010) and may influence therapeutic effects. Many previous ECS studies have utilized unstressed rodents, and though valuable to determine the effects of ECS, this approach falls short of understanding how ECT helps patients with depression. To appropriately model ECT, ECS must be paired with animals that show depressive-like behavior with similar neurobiological changes.

In the current study, we utilized the CSDS model in conjunction with a novel approach to ECS. Furthermore, ECS needs to better approximate the human ECT protocol. Currently, there are many differences, some involving rodent anatomy and electrode placement may not be reparable, but the anesthesia used, as it may impact recovery and neuronal plasticity, and the amount treatments per week, are modifiable. The present model addresses concerns regarding a lack of translationally relevant ECS which is typically delivered with isoflurane alone and daily administration for up to ten days. Thus, this new protocol has a higher likelihood of modeling human ECT. Our approach uses methohexital as an anesthetic as well as propofol administration upon recovery, as is the protocol for patients at the McLean Hospital ECT clinical service. Additionally, we administered ECS three times a week for two weeks, the typical frequency of ECT induction treatments. The current study sought to address prior limitations with mouse ECS models, validating behavioral responses as well as to begin to address molecular mechanisms of those responses. Thus, we examined the behavioral effects (social interaction and fear conditioning) after ECS in chronically stressed mice and measured bulk RNA transcription of the NAcc as it has been shown to mediate changes social reward in mice.

## Methods

### Animals

Wild-type C57/BL6 aged 5 weeks and CD1 retired breeders, aged less than 4 months obtained directly from Jackson laboratories and Charles River Laboratories, respectively, and were housed in a temperature-controlled room with free access to food and water and maintained on reverse 12hr/12hr dark/light cycle. Mice were habituated to light/dark cycle for at least 2 weeks prior to experiments. C57/BL6 were between 7 and 8 weeks at the time of social defeat. All behavioral procedures occurred during the dark cycle. All procedures were conducted in accordance with the guidelines established by the NIH and have been approved by the McLean Hospital IACUC.

### Drugs

10 mg/kg methohexital was made the day of each ECS session diluted with 0.9% saline and administered intraperitoneally (i.p., 45 mg/kg). Propofol (Zoetis Inc., Kalamazoo, MI) and administered intraperitoneally (i.p., 10 mg/kg). Isoflurane (Henry Schein, Dublin, OH) was administered nasally through a tube.

## Behavioral Procedures

### Chronic Social Defeat Stress (CSDS)

#### Materials

The materials used are described in Golden et al. (2014). Briefly, clear rectangular hamster cages (26.7 cm (w) x 48.3 cm (d) x 15.2 cm (h)) with a clear perforated Plexiglas divider (0.6 cm (w) x 45.7 cm (d) x 15.2 cm (h)) was used to house the aggressors and experimental and control animals.

#### Selection of Aggressors & Set-up

For the 6 days prior to CSDS, aggressors were screened for 5 minutes each day with a novel C57/BL6J mouse being placed in the aggressor’s home cage. Aggressive behavior and the total number of bites were noted. Aggressors selected bit within the first 5 seconds of screening and had at least one bite per minute. Aggressors were housed in new rat cages an hour after screening on the 6^th^ day and allowed to live in the full cage for 24hr with bedding squares. A divider was placed in the cage after the initial 24 hr with the bedding squares remaining on the aggressors’ side of the cage. All aggressors were on the same side of the divider and aggressors remained on their homecage side of the cage throughout the experiment. The intruder mice were never allowed to have bedding squares. CSDS started 24 hr after divider placement.

#### CSDS paradigm

*Control Condition*: For 10 consecutive days, C57/BL6J mice lived on either side of a Plexiglass divider in the same rat cages used for defeated animals. Every day, the mice were moved to a novel area (either to the other side of the same cage or a side of a different, novel cage), and allowed to briefly interact (∼5 seconds) with the control mouse that would be living on the other side of the divider for the next 24 hr. An hour after CSDS on the 10^th^ day, all control mice were single housed in standard mouse cages. *Defeated Condition*: For 10 consecutive days, intruder mice were placed in the homecage side (rat cage) of a novel aggressor for 5 minutes. After the 5 minutes, the intruders were placed on the other side of the Plexiglass divider. An hour after the defeats on the 10^th^ day, mice were single housed in standard mouse cages.

### Social Interaction Test (SIT)

#### Material

The field consisted of a square arena made of Plexiglass, measuring 42 cm x 42 cm x 42 cm. A wire-mesh enclosure measuring 13 cm (h) x 7.63 (w) x 7.63 (d) was placed in the center of the interaction zone. The interaction zone was an area 24 cm x 13.5 cm centered along one of the walls. The two corners were on the opposite wall, measuring 8.5 cm x 8.5 cm each, and mice were scored for time spent in these zones, with all four paws having to cross a dividing floor line to be considered an entry into any of these areas. All testing is conducted under reverse room lighting where behavior is continuously videotaped by a video camera placed over the structure and movement was tracked using EthoVision XT, version 8.5.

#### Selection of Aggressors

For the 2 days prior to the SIT, aggressors were screened for 5 minutes each day with a novel C57/BL6J mouse being placed in the aggressor’s home cage. Aggressive behavior and the total number of bites were noted. Aggressors selected bit within the first 5 seconds of screening and had at least one bite per minute. *The aggressors selected for the social interaction test must be novel to the C57/BL6J that will be tested*. If the SIT is occurring immediately after CSDS, the screening of aggressors for SIT will occur during days 9 and 10 of the CSDS paradigm. Two aggressors were selected for the SIT.

#### Social Interaction Test

Two trials were run for each mouse, each with duration of 150 seconds. In the first trial, there is not a target CD1 mouse in the wired mesh enclosure. In the second trial, there is a novel target CD1 mouse that had been previously screened in the metal barred cup, not permitting the target mouse to move, but still allowing for contact with the C57/BL6 mice. The two aggressor CD1 mice were alternated every 8 trials (every 4 C57/BL6 mice).

### Sucrose Preference

Preference for a 1% sucrose solution compared to water was examined (Maria João Primo, 2023). Mice were housed individually in standard shoe-box cages. To monitor water consumption, two polycarbonate water bottles were hung inside the cage (Animal Care Systems, Centennial, CO), filled with approximately 300 mL of water, and weighed. Bottle weights were recorded at approximately the same time daily. Mice drank from the two water bottles for 2-3 days to ensure that no strong side-bias was present for either bottle. When testing began, a 1% sucrose solution (Domino Superfine Quick Dissolve 100% Pure Cane Sugar, Yonkers, NY) was added to one of the bottles. One-half of each control and social defeated group received the sucrose bottle to the left and the other half was access to the sucrose bottle to the right of the bottle containing the water. Mice were presented with the two bottles, one with water and the other with 1% sucrose. The animals were allowed to drink freely for 24 h. The bottles were weighed prior to and after testing on day 1 and were returned to the cage with their positions counterbalanced for an additional 24 h. Preference for sucrose was calculated by subtracting the total water consumed from the total sucrose consumed and dividing this amount by the total fluid consumed. Positive scores indicate a preference for sucrose over water, negative scores denote a preference for water, and scores approaching zero indicate no preference.

### Auditory Threat Conditioning

Mice were habituated for two days and transported and maintained under dark light conditions. Then, on the third day, mice were fear conditioned with 5 trials of auditory CS (30s, 6kHz, 75dB) presentations paired with a shock US (1.0s, 0.7mA) with variable inter-trial intervals, administered in Context A, consisting of a darkened chamber equipped with a grid-floor for shock US administration. During fear conditioning, pre-CS, CS, and post-shock freezing were measured to examine fear acquisition. An automated program (FreezeFrame version 4, Coulbourne Instruments) was used to measure freezing, and an independent observer blind to all treatment conditions monitored accuracy. Fear memory consolidation was examined 24 hours later in a novel context (Context B), a darkened chamber equipped with a flat floor, which is critical for revealing deficits in auditory memory consolidation independent of contextual fear acquired to Context A with 30 trails of auditory CS (30s, 6kHz, 75dB) without a shock US. Freezing, defined as the lack of movement except what is necessary for respiration, was analyzed as the percentage of time spent freezing during the CS utilizing FreezeFrame. Fear memory extinction retention was then examined an additional 24 hours later in Context B, with 15 trials of auditory CS with the same parameters, again, without the shock US.

## Electroconvulsive Seizures (ECS)

Animals are weighed and received intraperitoneal injections of brevital at a dosage of 45 mg/kg. Upon loss of righting reflex, 2% isoflurane was administered until loss of toe-pinch withdrawal reflex. Then, animals were switched to 1% isoflurane. ECS Stimulation is then administered through the earclip electrodes coated in conductive electrode gel (identical as the product used in the McLean ECT clinic) through a pulse generator (Ugo Basile) at frequency of 100 Hz, pulse width of 5, shock duration of 2 seconds, and current of 80 mA. Subsequently, oxygen is administered until the return of the toe-withdrawal reflex. Then, mice are returned to home cages with handwarmers and an intraperitoneal injection of propofol at a dosage of 10 mg/kg and were responsive within 10 minutes. Sham animals were given methohexital intraperitoneal injections at a dosage of 45 mg/kg and at a loss of righting reflex put on 2% isoflurane for one minute, 1% isoflurane for an additional minute, and then subsequently switched to oxygen. There was no current conducted. Upon return of toe withdrawal reflex, they were given intraperitoneal injections of propofol at a dosage of 10 mg/kg and returned to their home cages with handwarmers.

## RNAseq Methods & Analysis

### Bulk RNA-seq Analysis

Total RNA was isolated using the Agilent absolutely RNA miniprep kit (Agilent Technologies, Palo Alto, CA, USA). Samples were processed by mRNA enrichment and rRNA removal. RNA sequencing libraries were prepared using the NEBNext Ultra RNA Library Prep Kit for Illumina following manufacturer’s instructions (NEB, Ipswich, MA, USA). Briefly, mRNAs were first enriched with Oligeno(dT) beads. Enriched mRNAs were fragmented for 15 minutes at 94 °C. First strand and second strand cDNAs were subsequently synthesized. cDNA fragments were end repaired and adenylated at 3’ends, and universal adapters were ligated to cDNA fragments, followed by index addition and library enrichment by limited-cycle PCR. The sequencing libraries were validated on the Agilent TapeStation (Agilent Technologies, Palo Alto, CA, USA), and quantified by using Qubit 2.0 Fluorometer (Invitrogen, Carlsbad, CA) as well as by quantitative PCR (KAPA Biosystems, Wilmington, MA, USA). The sequencing libraries were clustered on 1 lane of a flowcell. After clustering, the flowcell was loaded on the Illumina HiSeq instrument (4000 or equivalent) according to manufacturer’s instructions. The samples were sequenced using a 2×150bp Paired End (PE) configuration. Image analysis and base calling were conducted by the HiSeq Control Software (HCS). Raw sequence data (.bcl files) generated from Illumina HiSeq was converted into fastq files and de-multiplexed using Illumina’s bcl2fastq 2.17 software. One mismatch was allowed for index sequence identification.

### Data and Code Availability

RNA sequencing data is available at Gene Expression Omnibus (GEO) under GEO accession number: GSE234553 (A reviewer token can be obtained via the editor). The code for the analysis is freely available at https://github.com/klengellab/ECS_RNAseq_Maddox.

## RNA Sequencing

### RNA Sequencing Analysis

Fastq files were obtained from Genewiz by Azenta Life Sciences, South Plainfield, NJ and processed through the bcbio-nextgen (v 1.2.8) pipeline using the STAR aligner with mm39 reference genome (Chapman et al., 2021); Dobin et al., 2013; Mouse Genome Sequencing Consortium et al., 2002). After inspection of the quality control reports, Salmon quantified raw counts were used for downstream analysis in R 4.0.2 (Patro et al., 2017, R Core Team, 2021). Genes with counts less than 10 across samples were filtered and raw counts were normalized using variance stabilizing transformation (VST) in DESeq2 (v1.30.1). We used DESeq2 (v1.30.1) to identify differentially expressed genes between the sham and stimulation group (Love et al., 2014). SVA (v3.38.0) was used to calculate surrogate variables (SVs) to capture unwanted variance derived from unobserved factors such as cellular composition of the tissue sample (Leek et al., 2022). Inflation was adjusted with fdrtool (v1.2.15) and fdr corrected values were used to identify differentially expressed genes (DEGs) with a p_FDR_ < 0.05 (K, 2008).

### Gene Ontology (GO) Enrichment Analysis

All DEGs were used as input for the gene ontology enrichment analysis. We used clusterProfiler (v3.18.1) to detect fdr corrected significant gene ontology terms (p_FDR_ < 0.05) and org.Mm.eg.db (v.3.12) was used for annotation (Yu G, 2012), (M, 2019).

## Statistical Analysis

Statistical significance testing was determined by using two-way analyses of variance (ANOVA) and linear regressions. All statistical significance testing occurred with GraphPad Prism version 8. Post-hoc tests of multiple comparisons were performed using unpaired students t-test. Significance was determined as a *p-value* of < 0.05.

## Results

### Chronic social defeat stress yields social avoidance and a depression-like phenotype

We established a susceptible phenotype using 10-day CSDS (Figure 1A). Twenty-four hours later, the mice underwent the social interaction test (SIT), first without a target novel CD1 and then with the CD1 mouse present (Figure 1). The SIT stratifies mice between “susceptible” and “resilient” based on the ratio of time spent with a novel CD1 and the time spent exploring an empty cage (>1=resilient, <1=susceptible) (Krishnan V, 2007). Throughout our investigations, we had a low number of resilient mice (Supplemental Figure 2) and therefore focused on susceptible and unstressed control mice for this study (Figure 1B, *P*=0.0028), which more appropriately compares unstressed controls (n=14) with those that have a depression-like phenotype. Susceptible mice (n=13) spent less time in the social interaction zone when the target CD1 mouse was present compared to the unstressed mice (Supplemental Figure 1A, *P*=0.0041). Additionally, susceptible mice made less frequent entries into the interaction zone (Supplemental Figure 1B, *P*=0.0235), but did not display any differences in latency to the interaction zone when the CD1was present (Supplemental Figure 1C, ANOVA: group by target= *F*(3,48)=2.25, *P*=0.094). Susceptible mice and control mice showed differences in distance traveled during the SIT [ANOVA: group by target interaction *F*(3,48)=4.93, *P*=0.0046. Post-hoc unpaired student’s t-tests showed no difference between groups under conditions without a target (*P*=0.095) but a significant difference between groups under conditions with a target (*P*=0.0024)] as well as a significant decrease in control mice with the social target present (*P* =0.0085, Supplemental Figure 1D).

**Figure 1.**
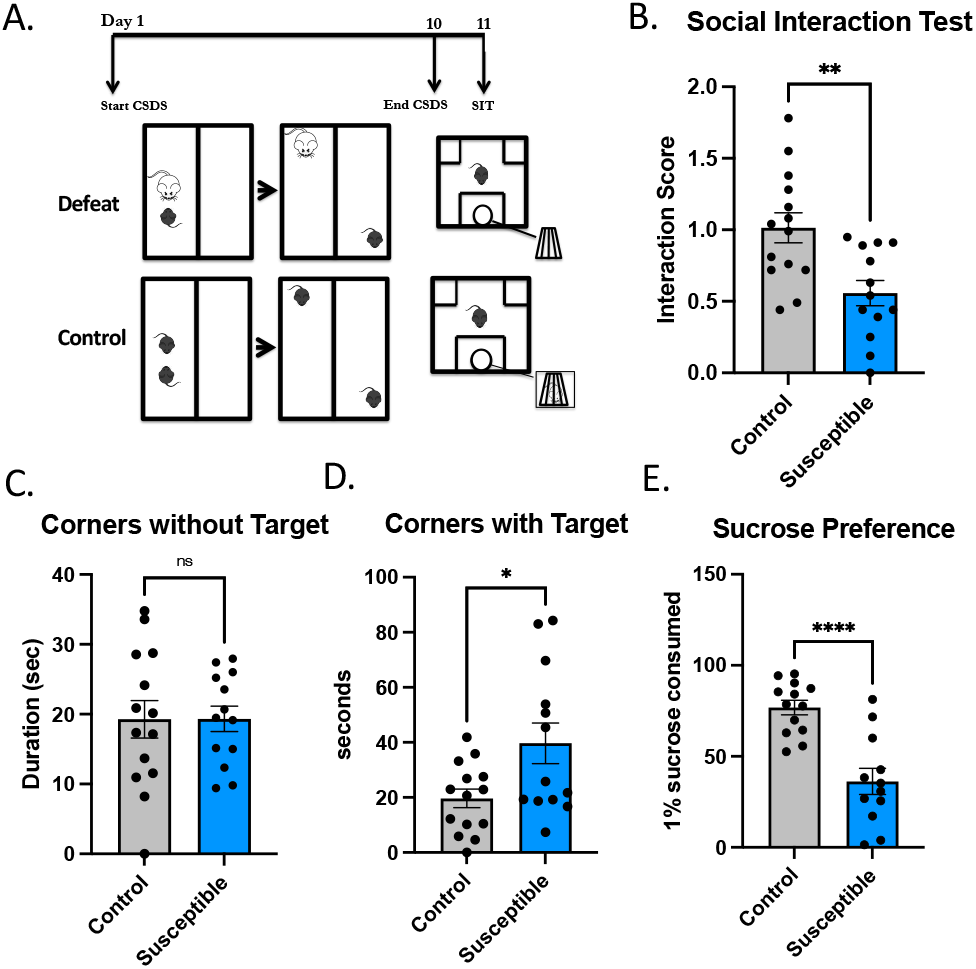
Mice subjected to chronic social defeat display social aversion and depressive-like behavior. **(A)** Diagram of the experimental design. **(B)** The presence of a social target decreased the amount of time susceptible mice (n=13) spent in the interaction zone (*P* = 0.0028) compared to control mice (n=14). There were no resilient mice in this cohort. **(C)** There was no significance difference in time spent in the corners between groups under no target conditions (*P* = 0.9856) but **(D)** there was a significant between groups under conditions with a social target (*P =* 0.0184)]. **(E)** Susceptible mice consumed significantly (*P* < 0.0001) less sucrose in the sucrose preference test.

**Figure 2.**
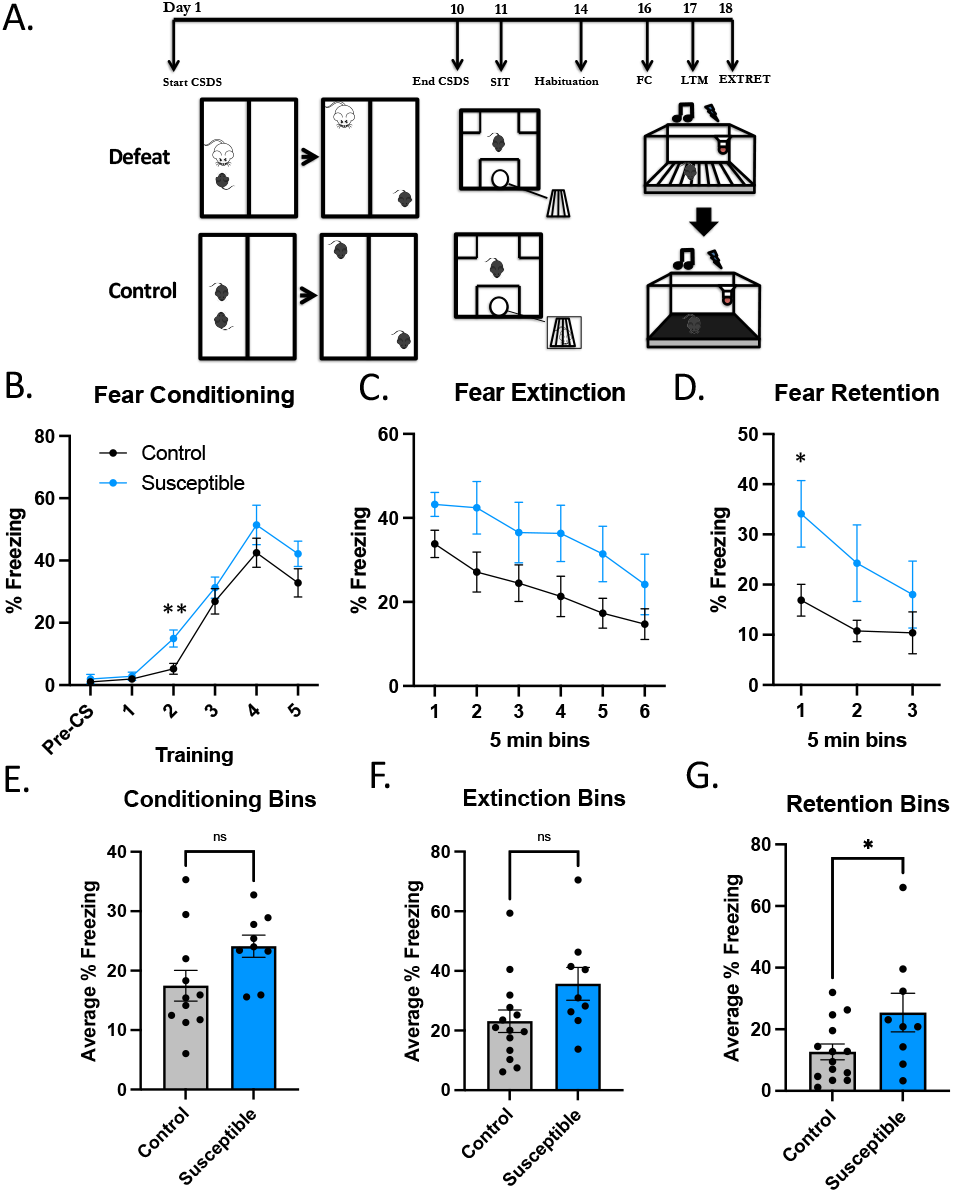
Chronic social defeat stress has little effect on threat-relevant memory extinction and retention. **(A)** Diagram of the experimental design. **(B)** Use of tone (US) and shock (CS) for CS-US pairings in the acquisition of fear memory. Percent freezing during fear conditioning over six five-minute bins showed no interaction [ANOVA: group by bins interaction F(5,126)=0.6944, P=0.6286], post-hoc pairwise comparisons showed a significant difference at timepoint 2 between control (n=11) and susceptible (n=9, P=0.0049). **(C)** Fear extinction over 30 minutes in six bins [ANOVA: group by bins interaction F(5,126)=0.1383, P=0.9831). **(D)** Retention of fear memories after extinction (extinction retention) had no group by trials interaction between control and susceptible mice [ANOVA: F(2,63)=0.4893, P=0.6154]; post-hoc student t-tests had one significant difference between control and susceptible mice at the first timepoint (P=0.01686). **(E)** Averaged fear conditioning bins of control and susceptible (P=0.0797). **(F)** There were also no differences in fear extinction between the groups (P=0.0641). **(G)** Retention of fear memories after extinction between control and susceptible mice was significantly different (P=0.0433).

During the SIT, in addition to exploring the CD1 mouse in the wire cup, mice are free to explore the arena, including the corners of the arena. Susceptible mice spent significantly more time in the corners when the CD1 was present (Figure 1D, *P*=0.018) than without the CD1 mouse (Figure 1C, *P*=0.99), compared to controls. Lastly, chronically stressed mice often display behavioral signs of reward and motivational dysfunction (Venzala E, 2012), (Russo SJ, 2013). To determine whether susceptible mice had a depression-like phenotype we employed the sucrose preference test (Figure 1E), which demonstrated a reduction in the amount of 1% sucrose consumed by susceptible mice compared to control mice (*P*<0.0001). The establishment of a susceptible and depression-like phenotype using CSDS enabled us to investigate behavioral and molecular changes by ECS.

### Chronic social defeat stress impairs fear extinction

Other investigators have demonstrated that CSDS has adverse effects on memory (C. Kyeremanteng, 2014). We set out to determine how susceptible mice respond to threat and how it impacts memory-related processes by employing auditory fear conditioning followed by extinction and retention. Auditory fear conditioning is a form of Pavlovian threat conditioning that pairs an unconditioned stimulus (US), in this case a tone, with a conditioned stimulus (CS), such as an electric foot shock, to elicit a conditioned response, such as immobility (freezing behavior). The amygdala is essential to freezing behavior, as seen in rodents with lesions to the lateral amygdala that exhibited altered freezing after threat conditioning (LeDoux JE, 1990). Due to the neuroplasticity, these acquired memories of threat and trauma can elicit heightened future responses that can be maladaptive in everyday life, relevant to the anxiety symptoms in depression and MDD’s high comorbidity with PTSD and anxiety disorders (Kessler RC, 1999), (JM, 1996), (Flory JD, 2015). Furthermore, these same brain regions are critical for recovery and extinction of fear memories, and anxiety- and trauma-related disorders have been repeatedly associated with extinction deficits.

After CSDS, mice were habituated to fear conditioning chambers for two days, then underwent auditory fear conditioning. We allowed 24 hours for fear memory consolidation then tested the mice for freezing in a novel context and underwent extinction trials. The following day, mice went through extinction retention testing. Fear extinction in susceptible (n=9) and control mice (n=11) had no significant overall difference (ANOVA: group by bins interaction *F*(5,126)=0.69, *P*=0.63), but there was a trend indicating possible increased fear consolidation and impaired extinction (Figure 2B-D) after social stress, as seen recently, though with modified protocols (Yang, 2023), (Goodman EJ, 2024). There was no difference (*P*=0.08) in mean percentage freezing over the six bins in control (*M=*17.47, *SEM*=2.57) and susceptible mice (*M*=24.11, *SEM*=1.88). Twenty-four hours after fear extinction, the same mice were subjected to extinction retention trials, where the tone (US) did not elicit a shock (CS). A group by trials repeated measures ANOVA revealed no significance [ANOVA: group by bins interaction *F*(5,126)=0.14, *P*=0.98)]. There were also no significant differences observed in mean percentage freezing over the six bins in control and susceptible mice during fear conditioning (Figure 2E) or extinguishing (Figure 2F), but there was a difference in extinction retention (Figure 2G, *P*=0.043), which can be interpreted as susceptible mice have an impairment in extinction.

Supplemental Figure 2 shows fear conditioning (Supp Fig 2A), extinction (Supp Fig 2B), and extinction retention (Supp Fig 2C) stratified by SIT ratio-determined resilient and susceptible, along with unstressed controls (Supp Fig 2A-G). The susceptible mice had the most differences from the resilient mice in fear conditioning (Supp Fig 2B) [ANOVA: group by bins interaction *F*(10,115)=2.61, *P*=0.0068]. Post-hoc unpaired comparisons showed significant differences at timepoints 3-5 between susceptible and resilient (*P*=0.0037, 0.012, 0.028, respectively), at timepoint 2 between control and susceptible (*P*=0.0049) and at timepoint 4 between control and resilient mice (*P*=0.023). When the fear conditioning bins were averaged, resilient mice had significantly lower freezing compared to susceptible mice [ANOVA: group by bins interaction *F*(2,23)=5.46, *P*=0.012); post-hoc students t-test showed a significant difference between susceptible and resilient mice (*P*=0.0024)]. There were no differences in extinction between all groups, but there was a significant increase in freezing behavior in susceptible mice compared to control mice [ANOVA: group by bins interaction *F*(2, 23) = 3.850, *P*=0.036; post-hoc unpaired students t-tests showed a significant difference between control and susceptible mice (*P*=0.043). This reiterates the previous finding of reduced extinction in susceptible mice compared to unstressed control mice. We then compared the same data collapsed into “defeated” (all mice that underwent CSDS, i.e., resilient plus susceptible) and unstressed controls (Supp Fig 2H-M), which, as expected reduced differences as it introduced additional variability.

### Chronic electroconvulsive stimulation reverses social avoidance behaviors following chronic social defeat stress

Electroconvulsive stimulation has been shown to reverse stress-induced phenotypes after chronic mild stress (Henningsen K, 2013) and exogenous cortisol stress (Hellsten J, 2002), but results from CSDS have not been consistent (van Buel EM, 2017). Therefore, we took susceptible and unstressed control mice and performed six sessions of ECS over two weeks (Figure 3A), the cadence typically seen when treating human depression during an ECT index course. One week following the sixth ECS session, the four groups of animals underwent the SIT to determine the effect of ECS on social aversion. There were no differences between control mice that underwent ECS compared to control sham mice (Figure 4B-H). Figure 3B shows our finding that susceptible mice that received ECS (n=7) had a higher SIT ratio than susceptible mice that received sham (n=6), ANOVA: group x treatment interaction *F*(3,21)=7.85; (*P=*0.011). Post-hoc unpaired students t-tests showed significant differences between susceptible sham and control sham (*P*<0.0001) and susceptible sham and control ECS (*P*=0.0024). Control mice that received sham ECS (n=7) also had a higher SIT ratio (approached social target more than the empty wire cup) than susceptible mice that received sham (n=6; *P*<0.0001). Susceptible mice that received ECS spent more time in the social interaction zone when the CD1 was present than susceptible mice that received sham (Figure 3D, *P*=0.018). Control mice that received sham treatment spent more time in the social interaction zone than susceptible mice that also received sham (Figure 3D, *P* <0.0001). We observed an increase in locomotion in susceptible mice that received ECS compared to sham (Figure 3E). Lastly, ECS significantly reduced the time susceptible mice spent in the corners during the SIT compared to sham when the social target is present (Figure 3H, *P*=0.0007), showing a reversal of anxiety-like behavior that was not present in control ECS (*P*<0.001) or sham mice (*P*<0.001). Supplemental Figure 3B shows ECS having a similar effect on defeated (all mice, resilient + susceptible) mice compared to defeated sham mice in the SIT (*P*=0.0039) and compared to control sham (*P*<0.0001). There were also differences in time spent in the interaction zone when the social mouse was present ANOVA: *F*(3,24)=13.47, *P<*0.0001. Post-hoc unpaired student’s t-test revealed significances between defeated and control sham (*P*=0.0002), control ECS and defeated sham (*P*<0.0001), defeated ECS and sham (*P*=0.0020), but not between control ECS and defeated ECS (*P*=0.062) or control sham and defeated ECS (*P*=0.68). Lastly, similar to the susceptible mice, there were significant differences in time spent in the corners when the social mouse was present, ANOVA: *F*(3,24)=16.20, *P<*0.0001. Post-hoc unpaired student’s tests revealed significant differences between sham control and defeated sham mice (*P*=0.0005), defeated sham and control ECS (*P*=0.0004) and defeated sham and ECS (*P*=0.0037), but not between susceptible and control ECS mice (*P*=0.11).

**Figure 3.**
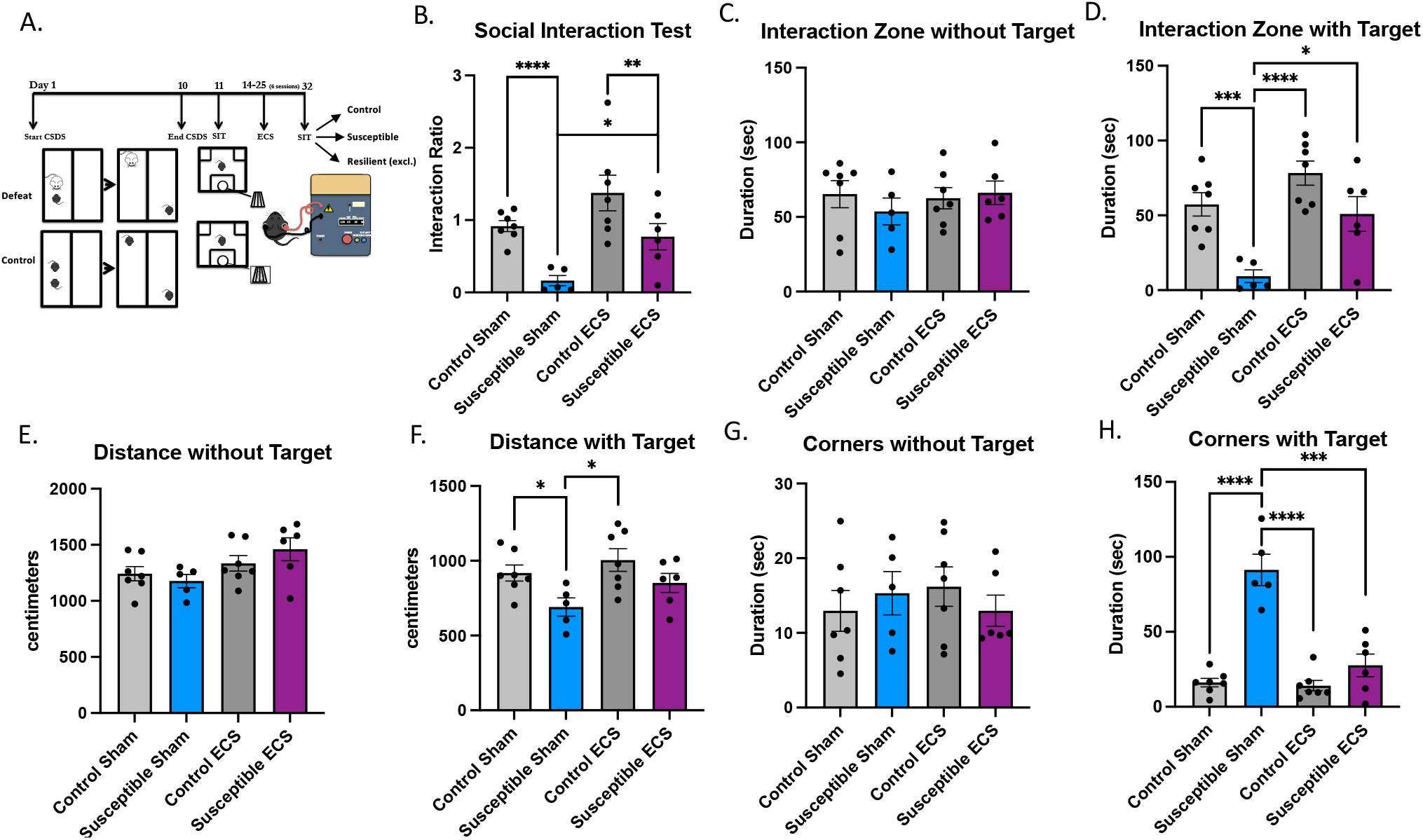
Reversal of social aversion from chronic social defeat stress after electroconvulsive stimulation. **(A)** Diagram of the experimental design. **(B)** The presence of a social target decreased the social interaction test of susceptible sham mice (n=5), which was not seen in susceptible mice (n=6) that received ECS (n=6); ANOVA: group x treatment interaction *F*(3,21)=7.851; (*P=*0.011). Post-hoc unpaired students t-test showed significant differences between susceptible sham and control sham (*P*<0.0001), susceptible sham and control ECS (*P*=0.0024), between susceptible sham and susceptible ECS (*P*=0.0178), but not control ECS (n=7) and susceptible ECS (*P*=0.0814) or control ECS and control sham (*P*=0.1015). **(C)** There was no significant interaction of time spent in the interaction zone between groups when the social target was absent (ANOVA: *F*(3,21)=0.4080, *P*=0.7488), but there were **(D)** significant differences between groups when the social target was present (ANOVA: *F*(3,21)=10.19, *P*=0.0002). Post-hoc unpaired student’s t-test revealed significances between susceptible and control sham (*P*=0.0007), control ECS and susceptible sham (*P*<0.0001), susceptible ECS and sham (P=0.0128), but not between control ECS and sham (*P*=0.0873) or susceptible ECS and control ECS mice (*P*=0.0732). **(E)** There were no significant differences in locomotion between the groups ANOVA: F(3,21)=2.400, P=0.0966, but there were when the social target was present **(F)** ANOVA: F(3,21)=3.822, *P*=0.0249. Post-hoc unpaired student’s tests revealed significant differences between sham control and susceptible mice (P=0.0196) and sham susceptible and control ECS (*P*=0.0126), but not between susceptible sham and ECS (*P*=0.1042). **(G)** There were no significant differences in the time spent in the corners of the arena when the social target is not present between the groups ANOVA: F(3,21)=0.4141, P=0.0745, but there were when the social target was present **(H)** ANOVA: F(3,21)=31.76, *P<*0.0001. Post-hoc unpaired student’s tests revealed significant differences between sham control and susceptible mice (P<0.0001) and sham susceptible and control ECS (*P*=0.0126), and between susceptible sham and ECS (*P*=0.0007).

**Figure 4.**
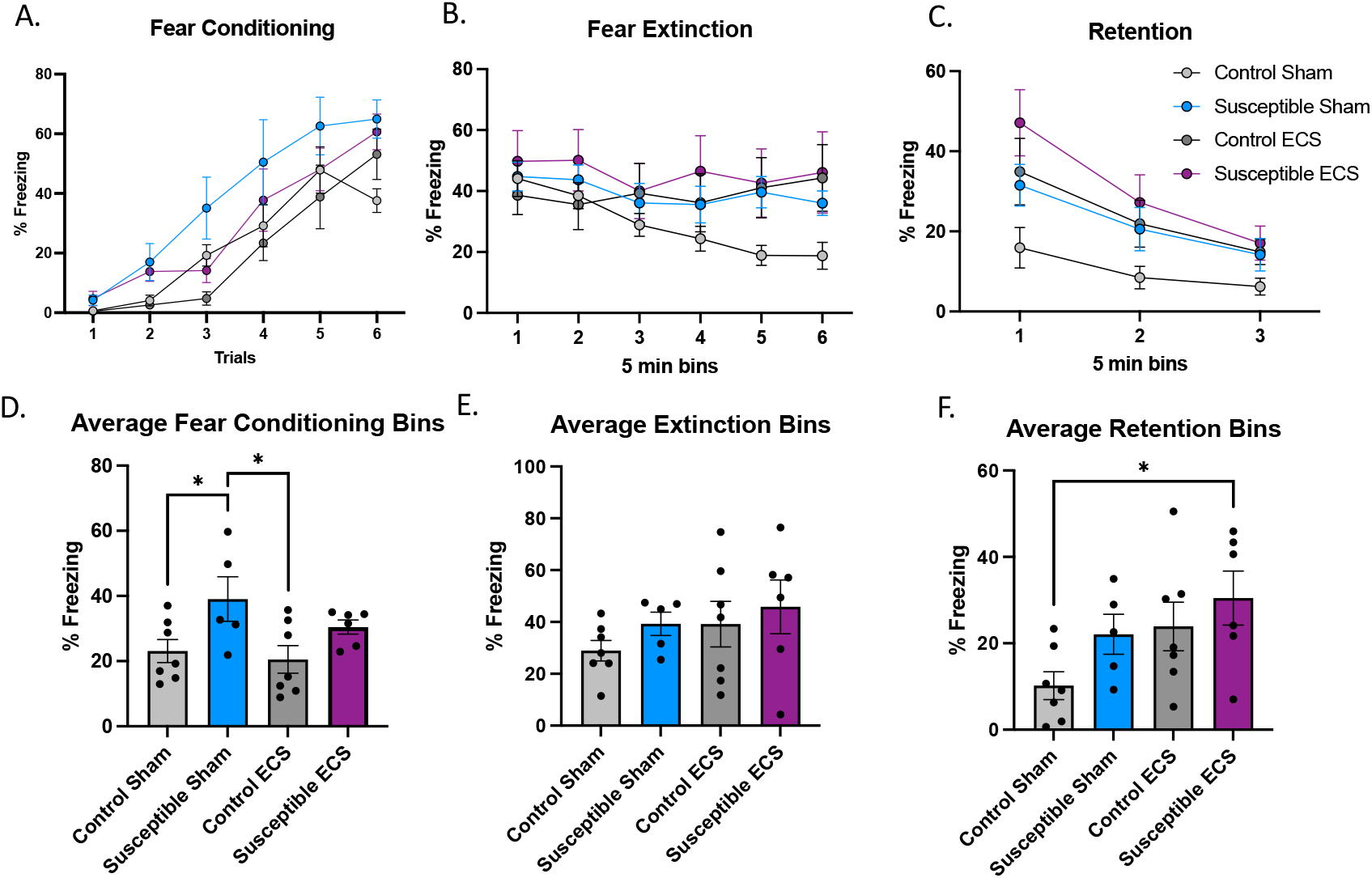
Electroconvulsive stimulation of mice after chronic social defeat stress enhances fear acquisition and retention. **(A)** Use of tone (US) and shock (CS) for CS-US pairings in the acquisition of fear memory in control mice or mice susceptible to CSDS and then either ECS or sham ECS. Percent freezing during fear conditioning over six five-minute bins showed no groups by trial interaction [ANOVA: F(15,126)=0.9153, *P*=0.5490]. **(B)** Fear extinction over 30 minutes in six bins [ANOVA: group by bins interaction F(5,126)=0.5243, P=0.9231) **(C)** Retention of fear memories after extinction (extinction retention) had no group by trials interaction between control and susceptible mice, ANOVA: F(2,63)=0.6173, *P*=0.7157. **(D)** Averaged fear conditioning bins of control and susceptible mice showed a significant group by treatment interaction ANOVA: F(3,21)=3.552, *P*=0.0319. Post-hoc unpaired students t-tests revealed significant differences between control and susceptible sham (*P*=0.0478) and between susceptible sham and control ECS (*P*=0.0350), but not between control and susceptible ECS (P=0.0735), susceptible ECS and control sham (*P*=0.1183) and susceptible sham and susceptible ECS (*P*=0.2263). **(E)** There were no differences in fear extinction between the groups, ANOVA: F(3,21)=0.8969, *P*=0.4596. **(F)** Retention of fear memories after extinction between control and susceptible mice was not significantly different, ANOVA: group by treatment interaction F(3,21)=0.8969, P=0.0567.

### Electroconvulsive stimulation following chronic social defeat stresses potentiates threat memory formation and impairs threat memory extinction

Clinically, ECT is limited by its negative effects on episodic memory (Porter RJ, 2020). Unfortunately, retrograde experiential memory loss is challenging to evaluate clinically. Typical memory screening tools, such as the Montreal Cognitive Assessment (MoCA) fail to capture subjective memory loss. In fact, a patients MoCA score may show little impairment and may even improve after ECT (Luccarelli J, 2022). Some of this can be attributed to improvement in the cognitive symptoms of depression (Bosboom PR, 2006). Episodic memory in rodents can be assessed with fear-learning (Kunčická D, 2024). This behavioral paradigm utilizes aspects of threat valence that can be quantified and help understand episodic memory formation, consolidation, and extinction (Figure 4). We utilized auditory fear conditioning to determine the effects of chronic stress and ECS on fear memory on encoding, extinguishing, and retention.

CSDS increased fear acquisition during conditioning in susceptible sham mice (n=5) compared to control sham (*P*=0.048) and control ECS mice [(*P*=0.035), ANOVA: *F*(15,126)=0.92, *P*=0.55] when the bins were averaged (Figure 4D). Susceptible sham mice exhibited more freezing during fear conditioning than control mice (both ECS and sham), though ECS had no apparent effect on fear acquisition in susceptible mice (Figure 4D, *P*=0.27). There was also no significant difference in fear extinction between any of the four groups over time (Figure 4B) or when the 5-minute bins were averaged (Figure 5E).

**Figure 5.**
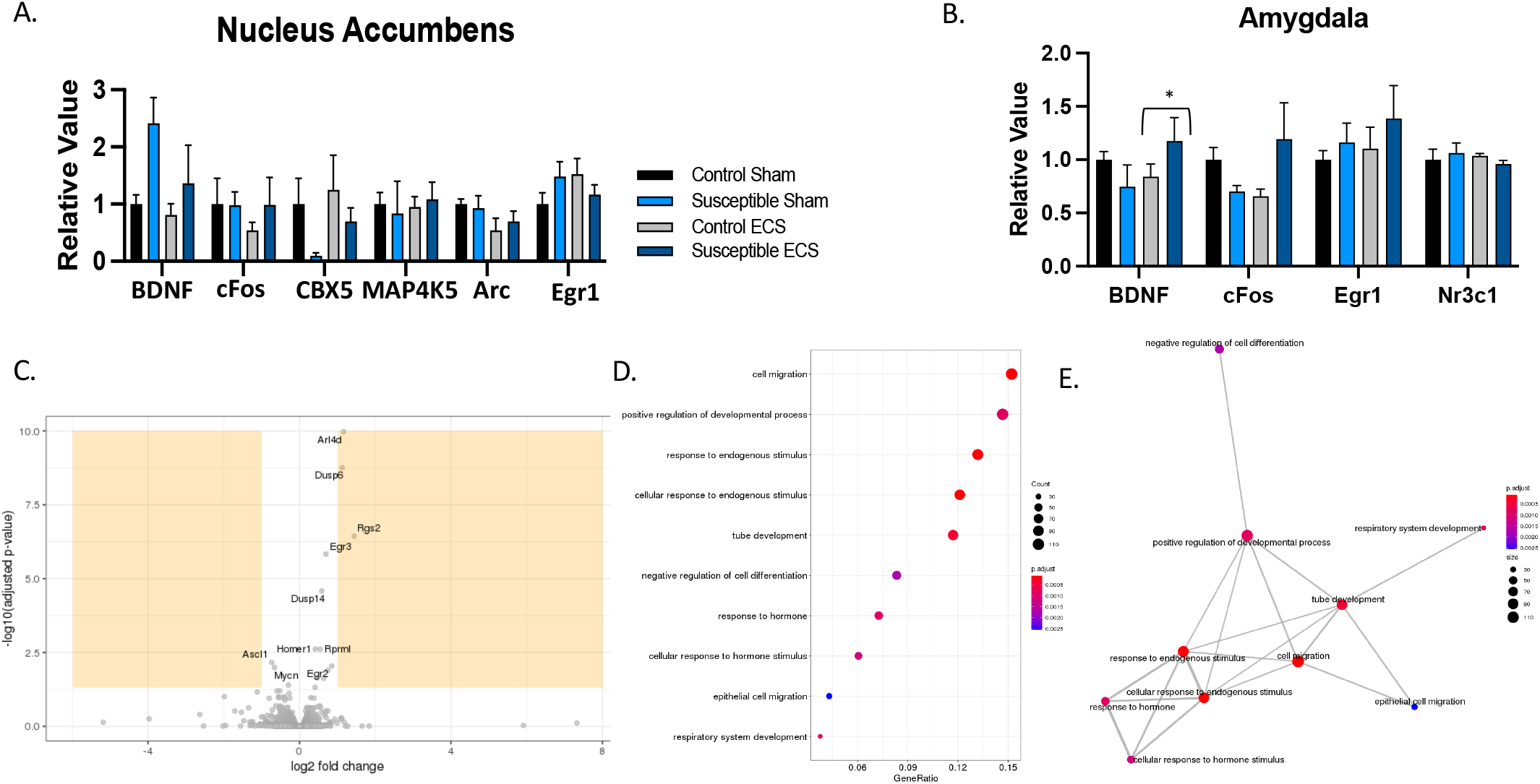
RNA levels in brain regions involved in stress and reward. **A)** RNA levels of BDNF, cFos, CBX5, MAP4K5, Arc, Egr1 from the Nucleus Accumbens **B)** RNA levels of BDNF, cFOS, Egr1, Nr3c1 after chronic social defeat stress and ECS/Sham with unstressed and sham controls, in the Amygdala. **C)** RNA sequencing from the Nucleus Accumbens revealed significant changes after ECS **D)** Gene Ontology Enrichment Analysis with nominal p-values E) Representation of cellular processes common within our RNA data using Gene Ontology Enrichment Analysis

When the resilient and susceptible mice are batched and analyzed as defeated mice (Supp Fig 4D), there is a significant group by treatment interaction (ANOVA: *F*(3,24)=3.23, *P*=0.041) in average fear conditioning of the five minute bins. Post-hoc unpaired students t-tests revealed significant differences between control and defeated sham (*P*=0.048) and between defeated sham and control ECS (*P*=0.026), implying that added resilient mice increased the power of the experiment (Supp Fig 4F, ANOVA: group by treatment interaction *F*(3,24)=3.214, *P*=0.041). Post-hoc unpaired student’s t-tests revealed significant differences between control and defeated sham (*P*=0.026), and between control sham and defeated ECS (*P*=0.0066).

### Chronic electroconvulsive stimulation regulates transcriptional processes in the Nucleus Accumbens and Amygdala

Since we are unable to readily determine molecular and cellular changes in humans undergoing ECT, mice offer an opportunity to understand gene expression in response to various stimulus. Therefore, we utilized quantitative real-time PCR (q-rtPCR) to determine changes in levels of plasticity-related RNA in brain regions shown to be involved in reward and stress in humans with depression. We assessed RNA levels in the NAcc and amygdala in mice that had CSDS and ECS, compared to controls (unstressed and sham). Figure 5 shows mRNA levels of plasticity-related genes in the NAcc (Figure 5A) and amygdala (Figure 5B). In the amygdala, chronic ECS significantly decreased cfos in unstressed mice (*P*=0.0285) but not in susceptible mice. There were no other differences in the transcription of these markers after CSDS and ECS.

Depression is often associated with social isolation and the NAcc has been shown to be involved in social reward learning in mice as well as anhedonia in humans (Russo SJ, 2013), (Panksepp JB, 2007), (Elmer, 2020). Considering our robust and consistent reversal of social avoidant behavior with ECS, we decided to further investigate ECS-dependent changes in the NAcc. We performed RNA-sequencing from the NAcc of unstressed mice that had 6 ECS or sham sessions (Figure 6C). ECS-induced RNA changes in the NAcc of unstressed mice showed increases in plasticity-related genes including HOMER1a, Rgs2, Egr3, Egr2, Sgk1, Arl4d, Dusp14, Rprml, Fhl2, Nr4a1, Akap5, C130074G19RiK) and 3 downregulated (Mycn, Ascl1, Tob1). Next, we performed a gene ontology enrichment analysis and found various cellular processes changing in response to chronic ECS (Figure 5), many involving cellular growth and neurogenesis, synaptic and cellular structure changes, in addition to regulators of calcineurin-NFAT signaling.

## Discussion

Depression is a debilitating mental illness that adversely affects millions of people. Depression affects the brain’s function by changing circuit, cellular, and molecular signaling. In stressed states, brains often react pathologically and demonstrate impairments in physiologic and appetitive processes, including reward. The reward system is particularly dysregulated in patients with depression, with anhedonia being a critical and common symptom. Both depression and ECT have been shown to adversely affect memory processes. We set out to determine phenotypic and molecular changes with chronic stress and a clinically relevant model of ECT to inform future investigations towards novel approaches for depression treatments.

Previous approaches implemented ECS on unstressed rodents, typically with up to ten daily treatments (Inta D, 2013), (van Buel EM, 2017), whereas patients usually receive ECT 2-3 times a week over the course of 2-3 weeks. In this study, we demonstrated that the social avoidance after CSDS can be reversed by ECS. This underscores the translational relevance of ECS using our approach. Observations in human neuroimaging led us to evaluate the NAcc and we found changes in expression of plasticity-related genes after ECS. Additionally, as seen in humans (Maren S, 2016), fear acquisition was increased with CSDS and subsequent ECS impaired fear extinction, which may reflect a negative impact on memory processes. Future work may be inclined to use this approach for additional insights into the mechanisms of ECT.

The mesolimbic pathway consisting of VTA dopaminergic projections to the NAcc contribute greatly to the rodent reward system. It has been demonstrated that the hippocampus coordinates VTA firing with direct projections, and disruption of the hippocampus could lead to reduced dopaminergic firing, potentially resulting in anhedonia (Pittenger C, 2008), (Nestler EJ, 2006). Furthermore, chronically stressed mice show that BDNF in the VTA-NAcc pathway is involved in social avoidance and other behaviors associated with depression (Eisch AJ, 2003) and blocking BDNF in this pathway reverses defeat-induced social avoidance (Berton O, 2006). Lastly, stressed mice display more excitatory postsynaptic currents in the NAcc, which may reflect decreased inhibitory neuronal activity (Christoffel DJ, 2011). The data presented in our study begins to illustrate mechanisms behind social aversion and link this behavior with VTA and NAcc circuit dysregulation, also observed in human depression.

We found that CSDS impaired fear memory extinction after fear learning. The amygdala and the PFC are implicated in fear extinction learning. Changes in the PFC have already been established in the CSDS model (Abe R, 2019), as well as seen in patients with depression (Pizzagalli DA, 2022), and alterations in the PFC-amygdala pathway could explain our observation of impaired extinction. CSDS has previously been examined for its ability to produce anxiety-like behaviors (Jianhua F, 2017), though some work has been done regarding direct alteration in amygdala-dependent memory formation in social stress (Goodman EJ, 2024).

The reversal of social avoidance in susceptible mice that received ECS can be the result of two possibilities. The first is that ECS changed emotional valence imputed by the VTA-NAcc circuit to reverse social avoidance behavior as ECS mice spent more time in the social interaction zone. The second possibility is that ECS induced social memory impairment. Susceptible mice that received ECS may have had cognitive impairment that increased social interaction because they may have forgotten the prior social stress they endured. This would necessitate “forgetting” many aspects of the experience, including resident attacks, resident odor, and other sensory input from the arena during the defeat, which would require much deeper investigation to disentangle.

Finally, the Ressler lab previously demonstrated projection-specific sequencing of populations in the prefrontal cortex and amygdala (e.g., McCullough et al, 2018). We observed ECS-induced RNA changes in the NAcc of unstressed mice that showed an increase in plasticity-related genes including Rgs2, Egr3, Egr2, HOMER1a, among others, and three that were downregulated (Mycn, Ascl1, Tob1). HOMER1a is a synaptic scaffold protein involved in calcium regulation, synaptic receptor trafficking, and facilitating MAPK signaling cascades. Rgs2 was shown to be integral to susceptibility phenotype in female mice; and EGR transcription factors, EGR3 in particular, mediate neurobiological adaptations to stress, reward, and social behaviors. Additionally, EGR molecules are involved in cytoskeletal regulation, including growth and stabilization of dendritic plasticity. Pharmacological targeting of the NMDA- and/or GABA-receptor trafficking processes through the modulation of these molecular processes after ECS may generate novel therapeutic approaches to influence the neuroplasticity for the treatment of depression and other stress-related disorders.

## Limitations

Though we modeled our protocol after human ECT, our ECS sessions occurred in an absence of a paralytic. In ECT, typically, succinylcholine is used as a paralytic agent, though succinylcholine cannot be used in mice because it is lethal upon injection. A situation that must be considered thereby ensuring that molecular changes are not occurring because of the motor function induced during seizures in ECS, but instead actually due to the brain-specific alterations that it elicits. To ensure measured molecular changes are only a response to ECS, as well as improve our model’s translational relevance, a different paralytic agent should be used in future studies. Additionally, it is important to note the use of propofol, an anesthetic that targets NMDA and GABA receptors, in our model of ECS. This drug is not commonly used following ECT, but McLean Hospital utilizes it because it has been shown to help patients reorient after anesthesia (Sakamoto A, 1999). It has been studied as a possible treatment for TRD (Mickey BJ, 2018), and has been examined for its role in improving cognitive function when administered with ECS in rats (Zhang F, 2019). Our dosage of propofol was likely too low to have these antidepressant effects, and defeated sham mice did not display behavioral improvements, thus our results are likely a response to ECS itself.

Most importantly, a major limitation of the CSDS model cannot be understated. A necessary consideration in studying neuropsychiatric conditions is the sexual dimorphism that exists in depression. Most individuals with depression are female, but most rodent studies of depression-like behaviors are conducted in males because female social defeat models are not well established. A strength of the male CSDS model is that it is founded on males’ natural territorial behavior, whereas both of the female defeat models require manipulations (e.g. chemogenetic alterations and the artificial application of a male scent). Thus, an appropriate female chronic stress model is necessary for future investigations into mood disorder related behaviors.

## Conclusion

In summary, utilizing the CSDS paradigm, we identified two behaviors that are altered following chronic stress: social avoidance and fear learning. Mice susceptible to CSDS spent less time in the interaction zone than their control counterparts when a social target was present. Additionally, defeated mice demonstrated heightened fear response and impaired fear extinction following auditory fear conditioning. We then examined how our novel ECS approach affects these two behaviors following CSDS. Specifically, six sessions of ECS over two weeks reversed social avoidance in susceptible mice, and then produced further impairment of fear extinction in defeated and control mice. This illustrates that CSDS and ECS may be an appropriate model of social isolation and reward as well as memory processes. In fact, ECT is known to have adverse effects on memory, particularly episodic memory. Thus, our model of ECS can be utilized to further study the neurobiological substrates underlying ECT. In turn, we will continue to investigate the mechanisms of ECT’s efficacy for depression and cognitive impairments, hopefully leading to increased treatment options for patients suffering from depression.

