## Supplemental Information for "A Clinically Aligned Murine Model of Electroconvulsive Stimulation Reverses Social Aversion and Displays Fear Memory Impairment After Chronic Social Defeat Stress"

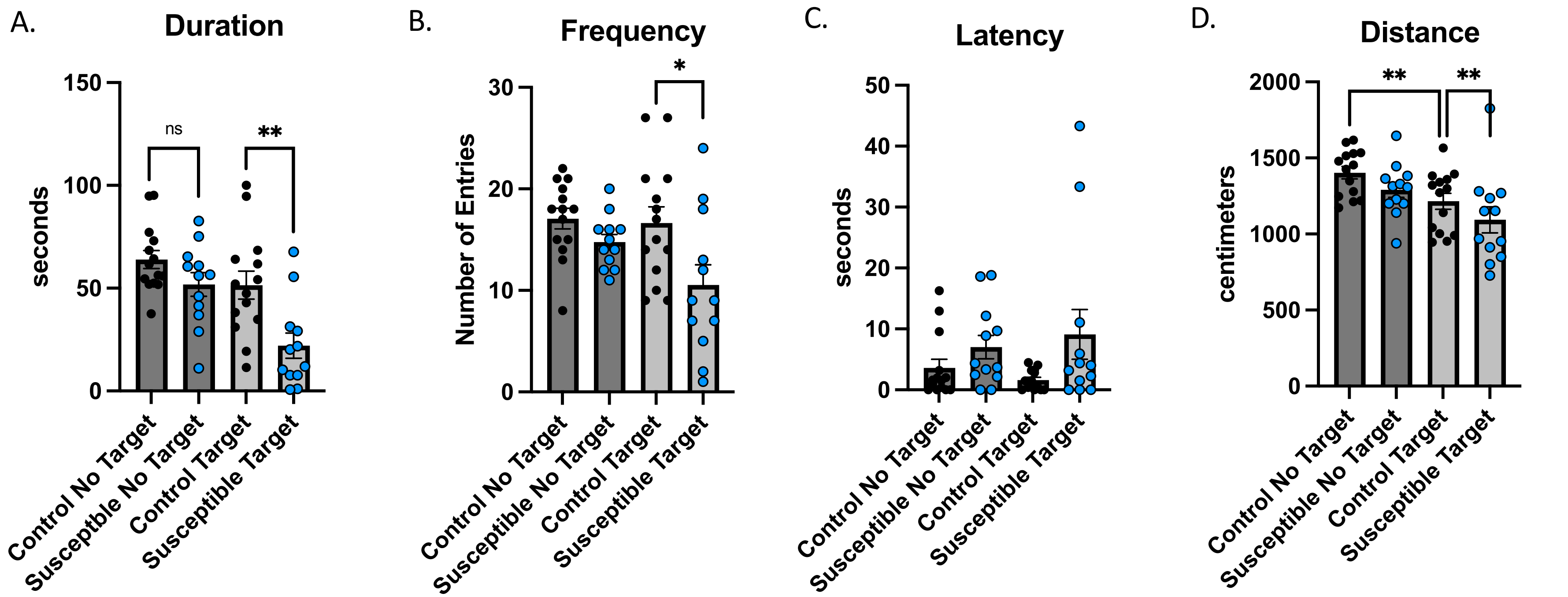


**Supplemental Figure 1.** Social interaction test details for susceptible and control mice. **(A)** The presence of a social target (CD1 mouse) decreased the amount of time susceptible mice (n=12) spent in the interaction zone compared to control mice (n=14, $P$=0.0041). The absence of a social target did not influence the time spent in the interaction zone. **(B)** The presence of a social target decreased the frequency of entries to the interaction zone in susceptible mice (n=12) compared to control mice (n=14) [ANOVA: group by target interaction $F$ (3,48) = 4.32, $P$ = 0.0089. Post-hoc students t-test between groups with the social target was significant (P=0.0235), while there was no difference without the social target (P=0.0874). **(C)** The presence of a social target did not change the time for the first entry latency into the interaction zone in susceptible mice (n=12) compared to control mice (n=14). [ANOVA: group by target interaction F (3, 48) = 2.253, $P$ = 0.0942. **(D)** Susceptible mice and control mice showed differences in distance traveled during the social interaction test [ANOVA: group by target interaction $F$(3,48) = 4.931, $P$ = 0.0046. Post-hoc student's t-tests showed no difference between groups under conditions without a target ($P$ = 0.0952) but a significant difference between groups under conditions with a target ($P$ = 0.0024)] as well as a significant decrease in control mice with the social target present (P=0.0085).

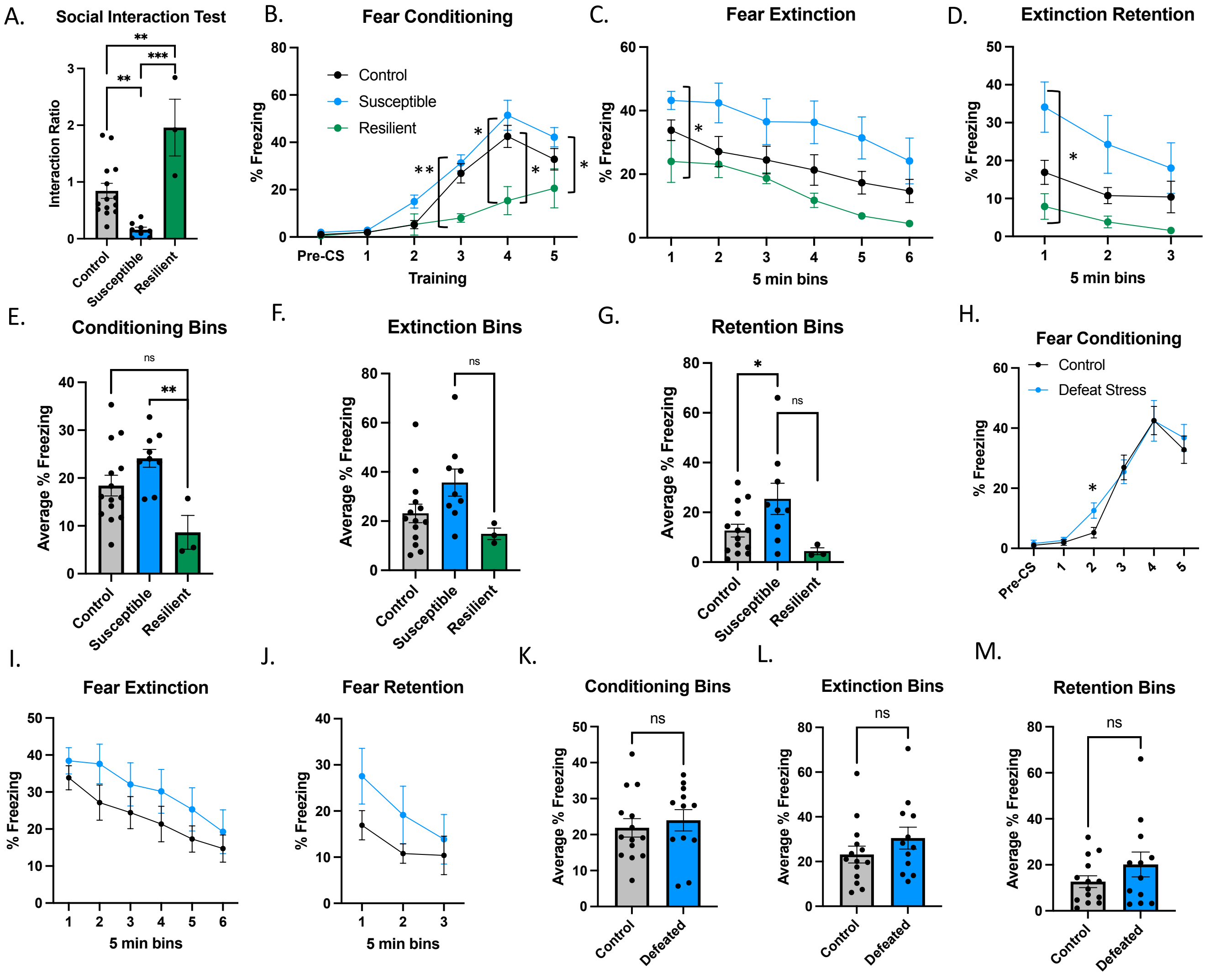


**Supplemental Figure 2.** Fear memory acquisition and extinction after chronic social defeat stress. **(A)** Mice displayed a resilient (n=3) phenotype defined by a SIT >1, while susceptible mice (n=9) and unstressed controls (n=14) [ANOVA: group by trials interaction $F(13,9)=8.6$, $P=0.0014$], post-hoc pairwise comparisons revealed significant differences between control and susceptible ($P=0.0011$), susceptible and resilient mice ($P=0.0001$) and between control and resilient mice ($P=0.0066$). **(B)** Use of tone (US) and shock (CS) for CS-US pairings in the acquisition of fear memory. Percent freezing during fear conditioning over six five-minute bins [ANOVA: group by bins interaction $F(10,115)=2.609$, $P=0.0068$], post-hoc unpaired comparisons showed a significant difference at timepoint 2 between control and susceptible ($P=0.00489$) and timepoints 3-5 between susceptible and resilient ($P=0.00373$, 0.01202, 0.02769, respectively) and at timepoint 4 between control and resilient mice ($P=0.02283$). **(C)** Fear extinction over 30 minutes in six bins [ANOVA: group by bins interaction $F(5,138)=3.562$, $P=0.0046$); post-hoc students t-test showed significant differences between susceptible and resilient mice at timepoint one ($P=0.0107$), and no significant differences between susceptible and control or between control and resilient mice. **(D)** Retention of fear memories after extinction (extinction retention) had no interaction between the three groups [ANOVA: group by trials interaction $F(4,69) = 0.2919$, $P=0.8823$]; post-hoc student t-tests had one difference between control and susceptible mice at the first timepoint ($P=0.0169$); there were no significant differences between control and resilient mice or susceptible and resilient mice. **(E)** Fear conditioning bins of control, susceptible and resilient averaged [ANOVA: group by bins interaction $F(2,23)=5.462$, $P=0.0115$); post-hoc students t-test showed a significant difference between susceptible and resilient mice ($P=0.0024$). **(F)** There were no differences in fear extinction between all three groups [ANOVA: group by bins interaction $F(2, 23) = 3.194$, $P=0.0597$. **(G)** Retention of fear memories after extinction between the three groups [ANOVA: group by bins interaction $F(2,23)=3.850$, $P=0.0361$; post-hoc students t-tests showed a significant difference between control and susceptible mice ($P=0.0433$). **(H-M)** Comparisons between control mice and all mice that underwent defeat stress (defeated = susceptible + resilient). **(H)** Control and defeated mice show no difference over CS-US pairings in the acquisition of fear memory [ANOVA: trials by group interaction $F(5,66)=0.6604$, $P=0.6547$. Post-hoc test showed no difference between groups under presentations of CS-US pairings except in trial 2, ($P=0.0238$)]. **(I)** 24 hours after fear conditioning, control mice (n=14) and defeated mice (n=12) underwent a series of fear extinction trials where the presentation of the tone (US) did not elicit a shock (CS). A group by trials repeated measured ANOVA revealed no significant differences, $F(5,144)=0.9868$. **(J)** 48 hours after fear extinction, control mice (n=14) and defeated mice (n=12) underwent a series of fear extinction trials where the presentation of the tone (US) did not elicit a shock (CS) to assess the retention of fear memories. A group by trials repeated measured ANOVA revealed no significant differences ($F(2,72)$, $P=0.7303$). **(K)** Fear conditioning bins of control and defeated mice were not significantly different with a students t-test ($P=0.5971$). **(L)** There were no differences in fear extinction between control and defeated mice ($P=0.2403$). **(M)** Retention of fear memories after extinction between control and defeated mice revealed no significant differences ($P=0.2013$).

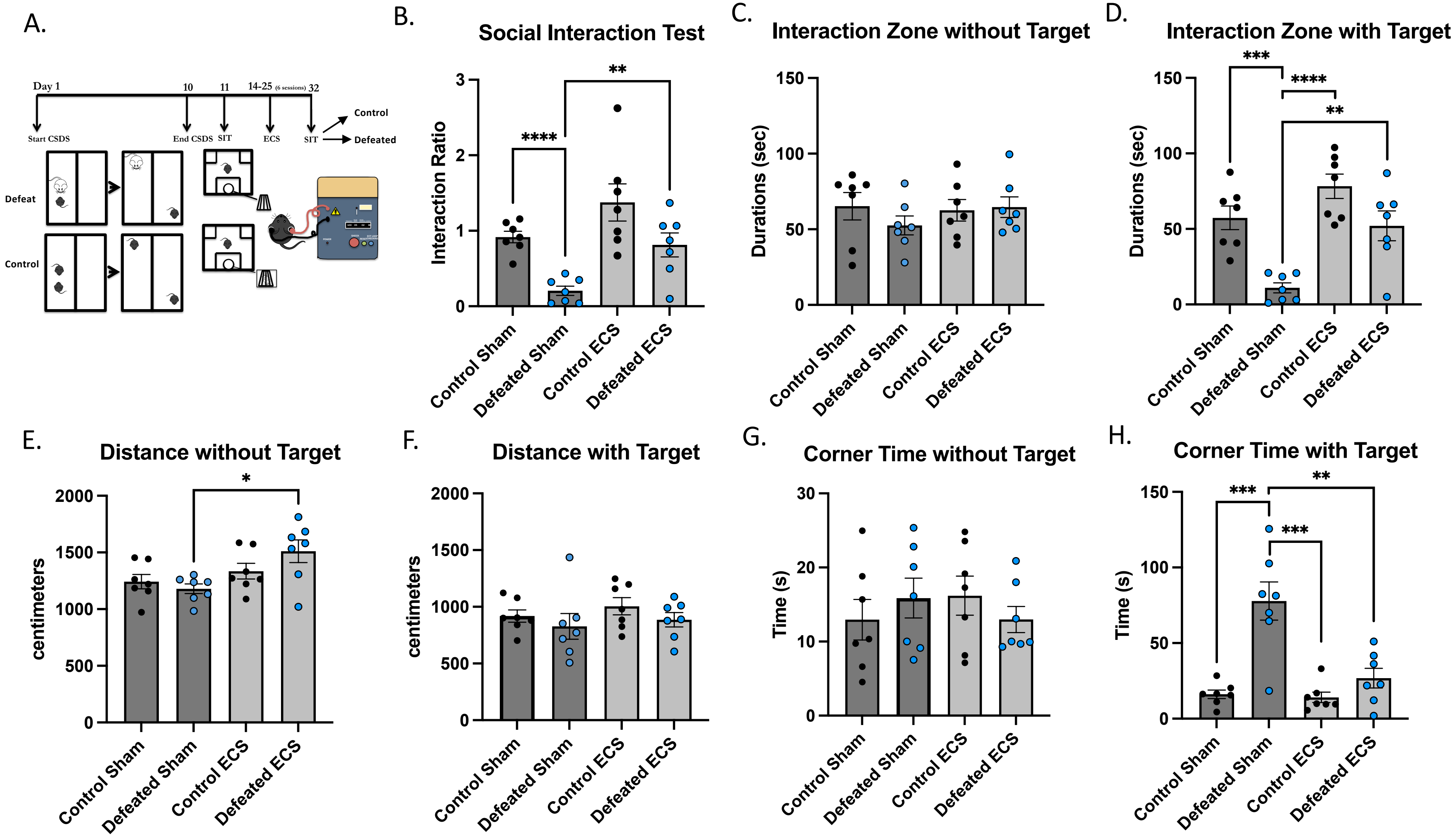
A.
Day 1
10
11
14-25 (6 sessions)
32
Start CSDS
End CSDS
SIT
ECS
SIT
Control
Defeated
Defeat
Control
B.
Social Interaction Test
Interaction Ratio
Control Sham
Defeated Sham
Control ECS
Defeated ECS
C.
Interaction Zone without Target
Durations (sec)
D.
Interaction Zone with Target
Durations (sec)
E.
Distance without Target
centimeters
F.
Distance with Target
centimeters
G.
Corner Time without Target
Time (s)
H.
Corner Time with Target
Time (s)

**Supplemental Figure 3.** Reversal of social aversion from chronic social defeat stress after electroconvulsive stimulation with resilient mice included. **(A)** Diagram of the experimental design. **(B)** The presence of a social target decreased the social interaction test of defeated sham mice (n=7) compared to control sham mice (n=7), ANOVA: group by treatment interaction $F(3,24)=9.683$, $P=0.0002$. Post-hoc unpaired students t-test revealed significant differences between control and defeated sham mice ($P<0.0001$), between defeated sham and defeated ECS ($P=0.0039$), but not between control sham and ECS (n=7), $P=0.1015$, or control sham and defeated ECS ($P=0.5642$), or control ECS (n=7) and defeated ECS ($P=0.0790$). **(C)** There was no significant interaction of time spent in the interaction zone between groups when the social target was absent (ANOVA: $F(3,24)=0.6428$, $P=0.5959$), but there were **(D)** significant differences between groups when the social target was present (ANOVA: $F(3,24)=13.47$, $P<0.0001$). Post-hoc unpaired student's t-test revealed significances between defeated and control sham ($P=0.0002$), control ECS and defeated sham ($P<0.0001$), defeated ECS and sham ($P=0.0020$), but not between control ECS and defeated ECS ($P=0.0622$) or control sham and defeated ECS ($P=0.6815$). **(E)** There was significant group by treatment interaction in locomotion between the groups ANOVA: F(3,24)=4.006, $P=0.0191$. A post-hoc unpaired students t-test showed a significant difference between defeated sham and ECS ($P=0.0103$). **(F)** There was not an interaction when the social target was present ANOVA: group by treatment $F(3,24)=0.8572$, $P=0.4767$. **(G)** There were no significant group by treatment interaction in the time spent in the corners of the arena when the social target is not present (ANOVA: F(3,24)=0.5068, $P=0.6813$), but there were when the social target was present **(H)** ANOVA: F(3,24)=16.20, $P<0.0001$. Post-hoc unpaired student's tests revealed significant differences between sham control and defeated mice ($P=0.0005$), defeated sham and control ECS ($P=0.0004$) and defeated sham and ECS ($P=0.0037$), but not between susceptible and control ECS mice ($P=0.1094$).

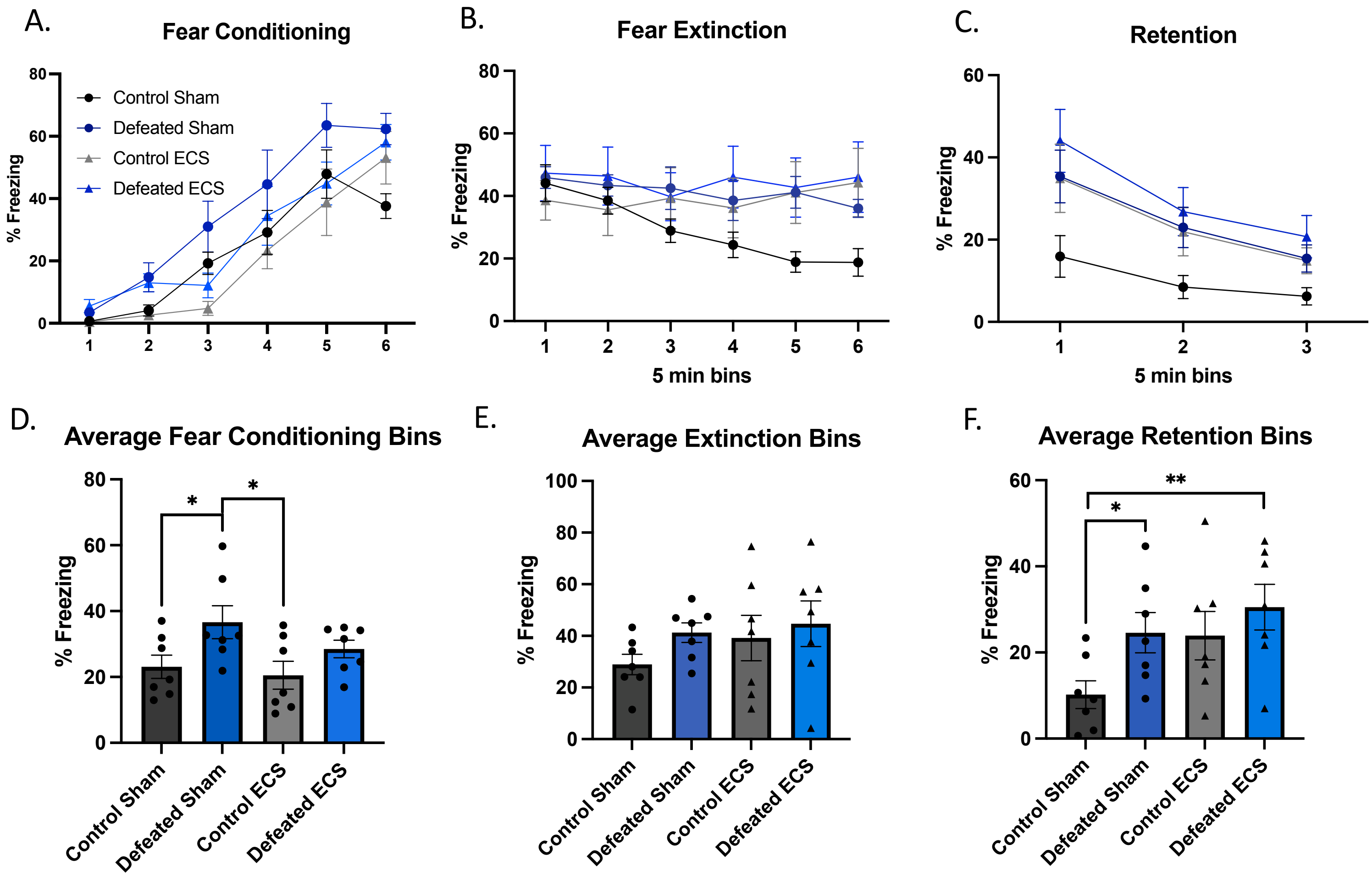


**Supplemental Figure 4.** Electroconvulsive stimulation of mice after chronic social defeat stress enhances fear acquisition and retention. **(A)** Use of tone (US) and shock (CS) for CS-US pairings in the acquisition of fear memory in control mice or defeated mice and then either ECS or sham ECS. Percent freezing during fear conditioning over six five-minute bins showed no groups by trial interaction, ANOVA: $F(15,144)=0.9965$, $P=0.4621$. **(B)** Fear extinction over 30 minutes in six bins [ANOVA: group by bins interaction $F(5,144)=0.7127$, $P=0.7687$) **(C)** Retention of fear memories after extinction (extinction retention) had no group by trials interaction between control and susceptible mice, ANOVA: $F(2,63)=0.6173$, $P=0.7157$. **(D)** Averaged fear conditioning bins of control and defeated mice showed a significant group by treatment interaction ANOVA: $F(3,24)=3.232$, $P=0.0401$. Post-hoc unpaired students t-tests revealed significant differences between control and defeated sham ($P=0.0476$) and between defeated sham and control ECS ($P=0.0302$), but not between control and defeated ECS (P=0.1363), defeated ECS and control sham ($P=0.2453$) and defeated sham and defeated ECS ($P=0.1788$). **(E)** There were no differences in fear extinction between the groups, ANOVA: $F(3,24)=0.9922$, $P=0.4132$. **(F)** Retention of fear memories after extinction between control and susceptible mice was significantly different, ANOVA: group by treatment interaction $F(3,24)=3.214$, $P=0.0408$. Post-hoc unpaired student's t-test revealed significant differences between control and defeated sham (P=0.0264), between control sham and defeated ECS (P=0.0066), but not between control sham and control ECS (P=0.0561), control ECS and defeated sham (P=0.9262), or defeated sham and ECS (P=0.4181).

Nominal p-values < 0.05 – 873 genes

| gene_id | baseMean | log2FoldChange | lfcSE | stat | pvalue | padj |
|---|---|---|---|---|---|---|
| Arl4d | 1007.50114 | 1.1621863 | 0.11631053 | 9.992099 | 1.650492e-23 | 1.155179e-19 |
| Dusp6 | 1641.05131 | 1.1276822 | 0.12558010 | 8.979785 | 2.712975e-19 | 9.494055e-16 |
| Rgs2 | 3202.49318 | 1.4450231 | 0.18190684 | 7.943754 | 1.961530e-15 | 4.576250e-12 |
| Egr3 | 3764.57729 | 0.6974991 | 0.09155288 | 7.618538 | 2.565651e-14 | 4.489248e-11 |
| Dusp14 | 1057.26382 | 0.5874298 | 0.08412976 | 6.982426 | 2.901260e-12 | 4.061184e-09 |
| Homer1 | 3922.49078 | 0.4193676 | 0.07165024 | 5.852983 | 4.828348e-09 | 4.827658e-06 |
| Rprml | 1266.20204 | 0.5465079 | 0.09315398 | 5.866716 | 4.445104e-09 | 4.827658e-06 |
| Ascl1 | 156.55670 | −0.7364989 | 0.13273699 | −5.548558 | 2.880357e-08 | 2.519952e-05 |
| Egr2 | 496.97254 | 0.8478173 | 0.15580421 | 5.441556 | 5.281721e-08 | 4.107418e-05 |
| Mycn | 167.11473 | −0.6598825 | 0.12256376 | −5.383994 | 7.285098e-08 | 5.098840e-05 |
| Sgk1 | 1898.18247 | 0.7443437 | 0.14056435 | 5.295395 | 1.187595e-07 | 7.556345e-05 |
| Fhl2 | 569.98997 | 0.4617027 | 0.09045690 | 5.104118 | 3.323411e-07 | 1.938379e-04 |
| Nr4a1 | 4162.48363 | 0.6366453 | 0.12549778 | 5.072961 | 3.916734e-07 | 2.108709e-04 |
| Tob1 | 895.35410 | −0.2918316 | 0.05960077 | −4.896440 | 9.758850e-07 | 4.878728e-04 |
| Akap5 | 3774.28524 | 0.4125044 | 0.08550957 | 4.824073 | 1.406561e-06 | 6.563014e-04 |
| C130074G19Rik | 644.07792 | −0.5981682 | 0.12696907 | −4.711133 | 2.463429e-06 | 1.061495e-03 |

Nominal P-value < 0.05; Log2fold change > 1

| gene_id | baseMean | log2FoldChange | lfcSE | stat | pvalue |
|---|---|---|---|---|---|
| Arl4d | 1007.501142 | 1.162186 | 0.1163105 | 9.992099 | 1.650492e-23 |
| Dusp6 | 1641.051311 | 1.127682 | 0.1255801 | 8.979785 | 2.712975e-19 |
| Rgs2 | 3202.493179 | 1.445023 | 0.1819068 | 7.943754 | 1.961530e-15 |
| Gm39741 | 14.467938 | 1.098928 | 0.3518999 | 3.122842 | 1.791141e-03 |
| Tbata | 13.920031 | 1.117258 | 0.3610578 | 3.094402 | 1.972099e-03 |
| Gm4926 | 10.905821 | 1.089456 | 0.3675822 | 2.963844 | 3.038223e-03 |
| Dct | 9.524389 | 1.182761 | 0.4522182 | 2.615465 | 8.910602e-03 |
| Gm18157 | 32.807271 | 7.328458 | 2.9732025 | 2.464837 | 1.370758e-02 |

Adjusted P-value < 0.05; Log2fold change > 1

| gene_id | baseMean | log2FoldChange | lfcSE | stat | pvalue |
|---|---|---|---|---|---|
| Arl4d | 1007.501 | 1.162186 | 0.1163105 | 9.992099 | 1.650492e-23 |
| Dusp6 | 1641.051 | 1.127682 | 0.1255801 | 8.979785 | 2.712975e-19 |
| Rgs2 | 3202.493 | 1.445023 | 0.1819068 | 7.943754 | 1.961530e-15 |

**Supplemental Figure 5.** Significant changes in bulk mRNA levels in Nacc ECS compared to Sham unstressed mice
